# Actions of camptothecin derivatives on larvae and adults of the arboviral vector *Aedes aegypti*

**DOI:** 10.1101/2021.09.06.458863

**Authors:** Frederick A. Partridge, Beth C. Poulton, Milly A.I. Lake, Rebecca A. Lees, Harry-Jack Mann, Gareth J. Lycett, David B Sattelle

**Author notes:** **Corresponding authors:** (DBS), (GJL). **These authors contributed equally.** (Frederick A. Partridge, Beth C. Poulton, Milly A.I. Lake).

## Abstract

Mosquito-borne viruses including dengue, Zika and Chikungunya viruses as well as parasites such as malaria and *Onchocerca volvulus* endanger health and economic security around the globe and emerging mosquito-borne pathogens have pandemic potential. However, the rapid spread of insecticide resistance threatens our ability to control mosquito vectors. Larvae of *Aedes aegypti* (New Orleans strain) were screened with the Medicines for Malaria Venture Pandemic Response Box, an open-source compound library, using INVAPP, an invertebrate automated phenotyping platform suited to high-throughput chemical screening of larval motility.

Of the 400 compounds screened, we identified rubitecan (a synthetic derivative of camptothecin) as a hit compound that significantly reduced *Ae. aegypti* larval motility compared to DMSO controls. Both rubitecan and camptothecin displayed concentration dependent reduction in larval motility with estimated EC50s of 25.5 ± 5.0 μM and 22.3 ± 5.4 μM respectively. We extended our investigation to adult mosquitoes and found that camptothecin increased lethality when delivered in a blood meal to *Ae. aegypti* adults at 100 μM and 10 μM and completely blocked egg laying when fed at 100 μM.

Camptothecin and its derivatives, inhibitors of topoisomerase I, have known activity against several agricultural pests and are also approved for the treatment of several cancers. Crucially, they can inhibit Zika virus replication in human cells, so there is potential for dual targeting of both the vector and an important arbovirus that it carries. Both humans and mosquitoes express the highly conserved topoisomerase I target, however, the design of derivatives with differing pharmacokinetic properties may offer a promising route towards the development of insect-specificity of this chemistry.

## Introduction

### Vector-borne diseases and pandemics

Humans have had to contend repeatedly with disease epidemics throughout history. Viruses such as Ebola, HIV, SARS-CoV-2 and Zika underscore the vulnerability of the human population to emerging pathogens. Furthermore, changes in our environment and society such as urbanisation, increased travel, and climate change will make epidemics more frequent and harder to control (Bedford et al., 2019). New and emerging infectious diseases, together with problems of anti-microbial resistance, are a challenge to our limited anti-infective medications and other tools for controlling diseases. To help to address this problem, the Medicines for Malaria Venture has recently launched the Pandemic Response Box, an open-source drug discovery program, where laboratories around the world collaborate by screening a library of structurally diverse compounds selected for potential activity against infective and neglected diseases.

Diseases transmitted by arthropod vectors endanger people in many areas of the globe. These vector-borne pathogens include protozoa, such as *Plasmodium, Trypanosoma* and *Leishmania*, nematodes, such as *Onchocerca volvulus,* as well as viruses, such as Chikungunya, Dengue, Yellow Fever and Zika (Shaw and Catteruccia, 2019). These diseases infect hundreds of millions of people, malaria kills 600,000 people each year and Dengue kills 40,000 (GBD 2017 Causes of Death Collaborators, 2018; GBD 2017 Disease and Injury Incidence and Prevalence Collaborators, 2018). The 2015 Zika epidemic, where the virus, spread via the mosquito *Aedes aegypti*, was introduced into the Americas and then spread rapidly, infecting perhaps 500,000 people, underscores vividly the pandemic potential of vector-borne diseases (Musso et al., 2019).

### Control of disease vectors

An important method for reducing the impact of vector-borne diseases is to target the vector. In the case of malaria, the incidence of clinical disease fell by 40% between 2000 and 2015, and it has been estimated that over half of this reduction was due to insecticide-treated nets (ITNs) that target the disease-transmitting *Anopheles* adult mosquitoes (Bhatt et al., 2015). However, ITNs have limitations, in particular the growing resistance to pyrethroids and other insecticides (Hemingway et al., 2016). ITNs are less useful for the control of pathogens spread by *Ae. aegypti,* which include chikungunya, dengue, yellow fever and Zika, as the mosquitoes prefer to feed outdoors at dawn and dusk. Larval source management is also important for vector control. This targets the larval stages of mosquitoes, which in the case of *Ae. aegypti* develop primarily in artificial, aquatic, urban environments, such as used tyres, drains, and sewers, with the aim of reducing the prevalence of the adult vector. Application of mosquito larvicides is an important component of larval source management. The major classes of larvicides in current use are chemical insecticides, typically organophosphates, such as temephos, the sodium channel targeting pyrethroids, bacterial larvicides such as *Bacillus thuringiensis* toxin, which impact gut cell membrane permeability, and insect growth regulators, like diflubenzuron, which block development of the insect.

However, resistance to current larvicides is a problem, with, for example, *Ae. aegypti* resistance to temephos, the major organophosphate larvicide, widespread in Brazil (Valle et al., 2019). And whilst pyrethroid use in water sources is now prohibited because of toxicity to fish (Farag et al., 2021), pyrethroids used in agriculture are known to leach into aquatic ecosystems. Even at the low concentrations observed, early larval exposure is thought to exacerbate the development of pyrethroid resistance in adults (Churcher et al., 2016) in areas where mosquito control is needed (Diabate et al., 2002) and that such larval stressors can impact the adult immune response (Hauser and Koella, 2020). Identification and development of new larvicidal agents thus remains a priority.

### Open science and the MMV Pandemic Response Box

Open science is an alternative way of doing science that aims to open up the research process, making innovation more efficient by the timely sharing of data, creation of collaborative communities and avoiding duplication of effort (Partridge et al., 2020; Todd, 2019). An example is the Medicines for Malaria Venture (MMV) Pandemic Response Box project, an open-source distributed drug-discovery project, where a compound library is screened in multiple laboratories in a diversity of assays. The goal is identification of small molecules with potential for development to control emerging diseases with pandemic potential. It follows on from the successful MMV Malaria and Pathogen Box projects (Van Voorhis et al., 2016; Veale, 2019).

We have developed a screening platform, INVAPP, that quantifies movement or growth of an organism in microplates (Partridge et al., 2018a). This system was originally developed to search for new anthelmintics (Hurst et al., 2014; Partridge et al., 2017, 2018b, 2021). We have recently adapted this platform for screening mosquito larvae of various species (Buckingham et al., 2021). Here we report the use of the INVAPP platform as part of the Medicines for Malaria Venture Pandemic Response Box project, by screening for new anti-mosquito compounds that could be useful in the control of vector-borne diseases.

## Materials and Methods

### Larval motility assay

*Ae. aegypti* egg papers were hatched in 500 ml pond salt solution (Blagon) supplemented with a quarter of a crushed 500 mg yeast tablet (Holland and Barrett), at 25 °C. After 18-24 h, larvae were collected with a 100 μm cell strainer, and diluted in pond salts solution to approximately 10 larvae per 100 μl.

Compounds were screened in 96 well plates. 100 μl of the larvae suspension was added to each well. For the primary screen using the Pandemic Response Box, the compound concentration in the assay was 10 μM, 1% v/v DMSO. Negative control (1% v/v DMSO) and positive control wells (10 μM deltamethrin, 1% v/v DMSO) were present on each assay plate. For the secondary screen, selected compounds were sourced from the original library material and screened at 10 μM, 1% v/v DMSO, with positive and negative controls as in the primary screen.

Movies were recorded and motility quantified using the INVAPP system (Buckingham et al., 2021; Partridge et al., 2018a). Movies of 200 frames at 100 ms intervals were recorded immediately after the larval suspension was pipetted into the assay plates (nominally 0 h timepoint) and again after 2 h and 24 h.

The Pandemic Response Box was a gift from the Medicines for Malaria Venture. For the primary screen, each well was normalised for inhomogeneity in the number of mosquitoes dispensed per well by dividing the motility score at 2 h or 24 h by that of the same well at 0 hours. The library was screened three times using independently prepared batches of mosquito larvae. Hit compounds were chosen where the median movement score at 2 h and/or 24 h was < 40% of the same wells at 0 h.

The secondary screen was carried out on two occasions, each time with five independent assay plates (n = 10). The 24-h time point was analysed. For each assay plate, the median movement score of the negative and positive control replicate wells was calculated and used for subsequent analysis. The effect of compound treatment was determined using a one-way ANOVA test, and the identity of active compounds was then determined by Dunnett’s test, in comparison with the DMSO negative control.

Rubitecan, and the related compounds camptothecin and topotecan, were then re-tested using solid material at an assay concentration of 100 μM. Camptothecin (208925), rubitecan (9-nitrocamptothecin, R3655) and topotecan hydrochloride (T2705) were obtained from Merck Life Science.

Concentration response curves were fitted using the R package *drc* (Ritz et al., 2015).

### Adult treatment assays

10 mM camptothecin stock was made in DMSO. Blood containing 100 μM and 10 μM camptothecin, 1% and 0.1% DMSO (as solvent controls respectively) and no additions (no DMSO control) was fed to 3 pools of 10 New Orleans adult (5-7 days old) females for each compound-concentration using a hemotek system. Adults were allowed 20 minutes to feed and any unfed adults were removed. Adults were maintained in paper cups supplied *ad libitum* with 10% sugar solution on cotton wool.

At 4 days post blood feeding, the surviving individuals (mortality recorded – 0-96 h) were transferred to a 5 mL bijou tube with a 2.5cm Whatman paper no.3 disk soaked in water pushed to the bottom to form a slight concave shape with a small pool of water for egg laying. Females were held in these tubes for 24 h to permit laying after which females were removed (mortality recorded - 96-120 h).

Lids were removed from tubes which were batch covered with netting to permit drying of the filter paper and eggs but preventing undesired egg laying by other mosquitoes. 7 days later 2 mL of yeast suspension (1 yeast tablet dissolved/suspended in 500 mL water) was added to each tube to stimulate larval hatching and netting was replaced. 2 days later the number of larvae hatched, and number of eggs laid were counted.

Differences in lethality, the number of eggs and larvae, and the percentage of laid eggs that hatched were assessed with a Tukey HSD post hoc test in R.

## Results

The actions on *Ae. aegypti* larval motility of each of the 400 compounds in the Pandemic Response Box was measured using the INVAPP system at 0, 2 and 24-hour timepoints. Fig 1A shows the effects of each compound tested, as well as DMSO-only and deltamethrin controls, on motility at 2h and 24 h. Deltamethrin at 10 μM effectively paralyses the larvae, but some compounds in the library showed some reduction in larval motility. The data for all 400 compounds in the MMV Pandemic Response Box are provided in the S1 Table. Fourteen compounds, highlighted in blue in Fig 1A, that reduced motility at 2h and/or 24 h to less than 40% of controls were selected as candidate hits and taken forward to a secondary screen. These compounds were retested at 10 μM in a secondary screen (Fig 1B), where the effects on motility after 24 h of treatment were analysed. A one-way ANOVA test found a significant effect of compound treatment on motility [F(15,144)=3.891 p=7.86e-06]. Dunnett’s test was then used to compare each compound with the DMSO-only control. Deltamethrin (the positive control, P = 0.027) and rubitecan (P = 0.041) showed a significant difference in motility compared to the control. The structure of rubitecan is shown in Fig 1C. Rubitecan is a topoisomerase inhibitor, originally developed as a potential therapy for various cancers (Clark, 2006).

**Figure 1.**
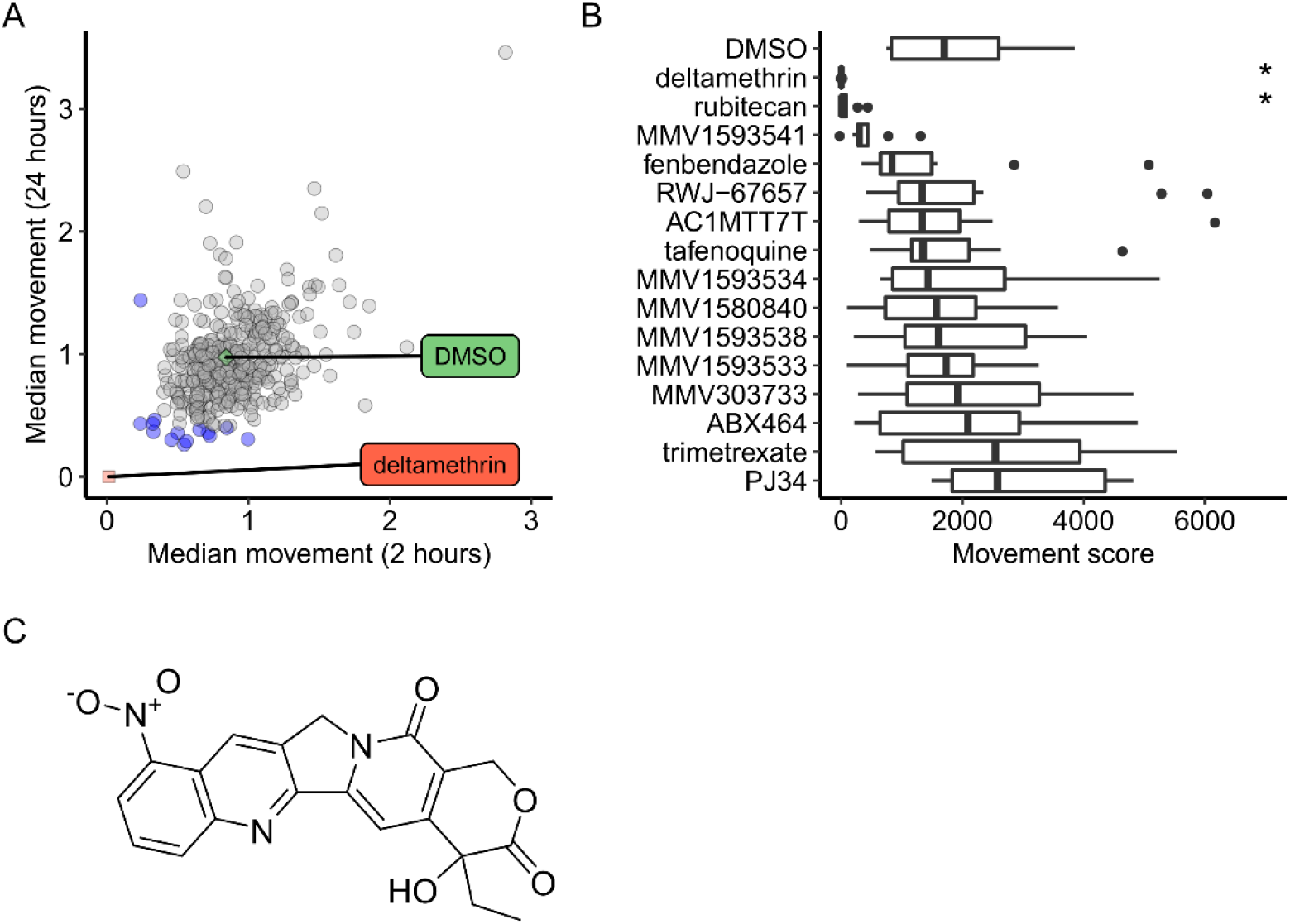
Screening the 400 compound MMV Pandemic Response Box chemical library in an *Ae. aegypti* larval motility assay led to the identification of the hit compound rubitecan. (A) Primary screen. Each point is the effect of one compound on motility at 2h and 24 h, normalised to the motility at the 0-h timepoint. n = 3. DMSO-only and deltamethrin were the negative and positive control compounds respectively. Blue points indicate the 14 compounds that were selected as candidate hit compounds. (B) Secondary screen, showing the effects of each compound on motility after 24 h. n=10. * indicates P < 0.05 compared to the DMSO-only control (Dunnett’s test). (C) Structure of rubitecan.

Having pursued these initial studies using library material stored as DMSO stocks, it was important to confirm the activity of rubitecan in the larval motility assay using solutions freshly prepared from solid material. Rubitecan is a synthetic derivative of camptothecin (Fig 2B), an alkaloid isolated from *Camptotheca acuminata*, a tree native to China. We also wanted to determine the activity of camptothecin itself, as well as topotecan (Fig 2C), another camptothecin derivative, which is approved for the treatment of cervical, ovarian and small cell lung cancers. These compounds were tested in the same 24-hour treatment larval motility assay at 100 μM. These results are shown in Fig 2A. A one-way ANOVA showed a significant effect of treatment, F(3,16)=22.0 p=6.32e-06. Dunnett’s test was then used to compare each treatment with the DMSO-only control. Camptothecin (P = 8.5e-6), rubitecan (P = 8.3e-6) and topotecan (P = 0.00087) all showed a significant difference in motility compared to the control, although the effect on motility was less in the case of topotecan. Examples of mosquito morphology and movement in wells treated with each compound are presented in the S1 movie. A time-lapse montage is also shown in Fig 2D.

**Fig. 2.**
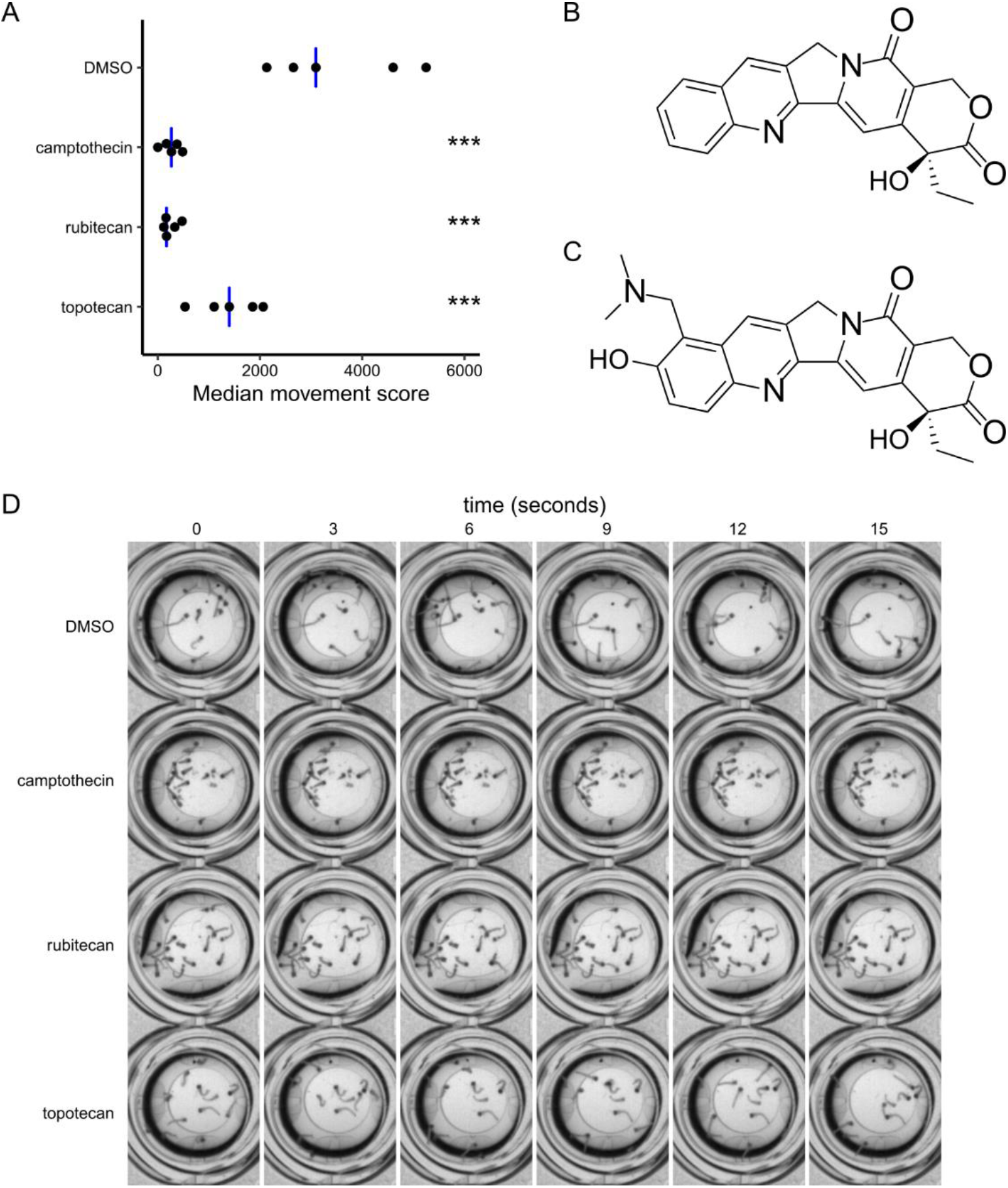
Re-testing the actions on larval motility of rubitecan and related compounds prepared freshly from solid material. (A) Retesting in the *Ae. aegypti* larval motility assay of rubitecan prepared from solid material and testing of two related compounds, camptothecin and topotecan, also prepared from solid material. Compounds were screened at 100 μM. Each point is the within-batch median movement score for each of n=5 biological replicates (batches of independently hatched mosquito larvae). Blue bar indicates the between-batch median. A one-way ANOVA showed a significant effect of compound treatment F(3,16)=22.0 p=6.32e-06. A post-hoc Dunnett’s test was then used to compare compound treatments with the DMSO-only control. *** indicates P < 0.001. (B) Structure of camptothecin. (C) Structure of topotecan. (D) Time-lapse montage of representative assay wells. This is presented as video in the S1 movie.

We next wanted to determine the concentration dependence of the larvicidal effect of the hit compounds. Concentration-response curves were obtained using the same larval motility assay (Fig 3A,B). Curves were fitted using the 4-factor log-logistic model. The EC_50_ of camptothecin was estimated to be 22.3 ± 5.4 μM, and that of rubitecan estimated to be 25.5 ± 5.0 μM.

**Fig. 3.**
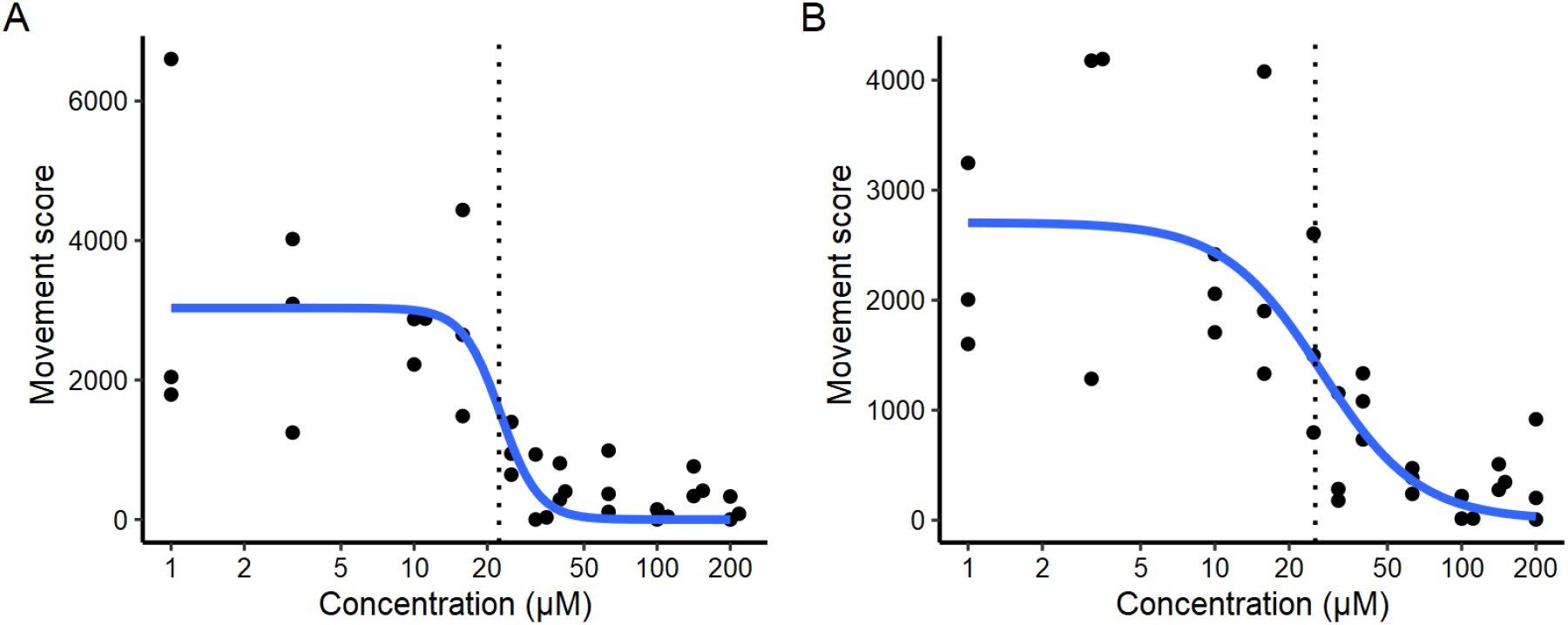
Concentration dependence of the actions on larval motility of (A) camptothecin and (B) rubitecan. Curve fitted using the 4-factor log-logistic model. n=3. Dotted lines indicate the EC_50_.

Finally, we investigated the usefulness of camptothecin to target adult mosquitoes. Attempts to kill adult mosquitoes in tarsal assays (100 μM, 30 min) and in sugar meals (100 μM, daily) did not indicate a strong phenotype (data not shown). Females were fed with blood containing camptothecin according to the regimen outlined in Fig 4A and mortality was recorded at each time point. The results are shown in Fig 4B and C. There was a significant difference in lethality between treatment groups as determined by one-way ANOVA (F(4,10) = 40.72, *p* = 3.7×10^−6^). A Tukey post hoc test (95% CI ± 25.7) revealed significant increased mortality in females fed 100μM camptothecin compared to those fed no compound (+80.0%, *p* = 9.8×10^−6^), 1 % DMSO (+76.7%, *p* = 1.45×10^−5^) and 10 μM camptothecin (+41.2%, *p* = 0.0026) and in those fed 10 μM camptothecin compared to those fed no compound (+38.8%, *p* = 0.004) and 0.01% DMSO (+39.4%, *p* = 0.0036).

**Fig 4.**
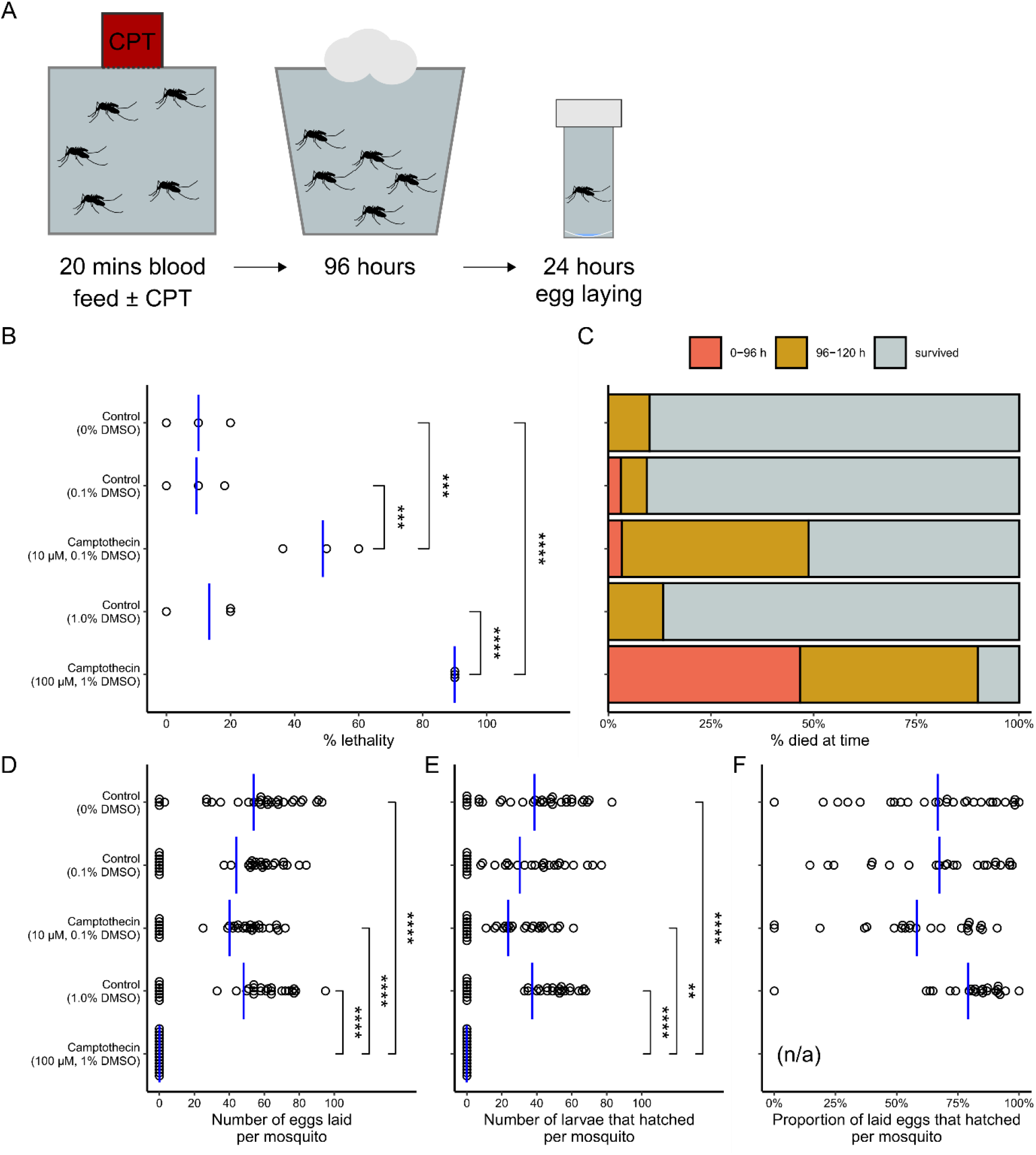
Assessing the effects of camptothecin on adult *Ae. Aegypti.* (A) Diagrammatic representation of the methodological process. (B) Effect of camptothecin concentration on percentage lethality across the entire experiment - each point represents the percentage lethality in each experimental replicate (n=3). (C) Effect of camptothecin concentration on percentage of total females which died indicating the time period during the experiment when death occurred (red = 0-96 h, yellow = 96-120 h) or survived to the end of the experiment (grey). (D) Effect of camptothecin concentration on the number of eggs laid per individual mosquito. Each point represents the number of eggs laid by an individual female (n = 30 per condition). (E) Effect of camptothecin concentration on the number of larvae hatched per individual mosquito. Each point represents the number of larvae hatched from an individual female (n = 30 per condition). (F) Effect of camptothecin concentration on hatch percentage. Each point represents the proportion of eggs which hatched for each individual female (n = 30 per condition). No females laid eggs and so no hatch percentage could be calculated where (n/a) is noted. Vertical blue lines indicate the mean and significance as determined using a Tukey post hoc assessment is indicated as follows on B, D, E and F (**** < 0.0001, *** < 0.001, ** < 0.01, * < 0.05, ‘absence of bracket’ > 0.05).

We also measured the number of eggs laid per adult mosquito (Fig 4D), the number of larvae that hatched per adult mosquito (Fig 4E), and the proportion of eggs that hatched for each treated adult mosquito (Fig 4F). Significant differences in the number of eggs laid were also observed between treatment groups using a one-way ANOVA (F(4,131) = 12.52, *p* = 1.16×10^−8^). Females exposed to 100 μM camptothecin did not lay any eggs. A Tukey post hoc (95% CI ± 22.19) indicated significant differences in the number of eggs laid by females fed 100 μM camptothecin and females fed no compound (− 53.9 eggs, *p* < 1×10^−7^), 1% DMSO (− 48.13 eggs, *p* = 2×10^−7^) and 10 μM camptothecin (− 40.1 eggs, *p* = 1.81 ×10^−5^).

The number of larvae that hatched also differed significantly by treatment using a one-way ANOVA (F(4,131) = 9.846, *p* = 5.38×10^−7^) but the significant reductions observed with a post hoc Tukey (95%CI ± 18.9) were between females fed 100 μM camptothecin and females fed no compound (− 38.7 larvae, *p* = 9×10^−7^), 1% DMSO (− 37.4 larvae, *p* = 2.3×10^−6^) and 10 μM camptothecin (− 23.7 larvae, *p* = 0.00632).

No significant differences in egg laying and number of larvae hatched were observed between females fed 10 μM camptothecin and controls and no significant differences in larval hatch percentage between treatments were detected using a one-way ANOVA (F(3,87) = 2.687, *p* = 0.0514).

## Discussion

### Camptothecin derivatives as insecticides

In this study, we screened the MMV Pandemic Response box in a mosquito larval motility assay, and identified that camptothecin, as well as the derivatives rubitecan and topotecan, had anti-larval activity against *Ae. aegypti.* This observation is concordant with previous observations of camptothecin-related compounds having insecticidal properties, although no compound from this chemotype has reached the market for this use (Liu et al., 2015). Indeed, a crude extract of *C. acuminata* was traditionally used in China to control pests (Zhang et al., 2012). Camptothecin was first shown to have chemo-sterilant activity against the housefly (DeMilo and Borkovec, 1974), and camptothecin or derivative compounds are active against agricultural pests (Liu et al., 2010a; Ma et al., 2010).

### Camptothecin derivatives as antivirals

Camptothecin derivatives, including topotecan and irinotecan, have been approved for the treatment of various cancers. They are inhibitors of topoisomerase I (TOP1), an enzyme important for DNA replication and repair, as well as transcription. Because of this mechanism, camptothecin derivatives have been investigated as potential antivirals (Pantazis et al., 1999). Of particular note, camptothecin or derivatives have shown activity in cells against herpes simplex virus type 2 (Liu et al., 2010b) and enterovirus 71 (Wu and Chu, 2017). Interestingly, camptothecin also suppresses the host response to viral and bacterial infection and protected mice in a model of lethal inflammation (Rialdi et al., 2016).

The dual insecticidal and anti-viral activities of camptothecin-like compounds is intriguing and may motivate further study. Zika virus is transmitted vertically within the *Ae. aegypti* population (Costa et al., 2018; Thangamani et al., 2016), and vertical transmission in the mosquito host also occurs with many other flaviviruses. Furthermore, *Ae. aegypti* larvae can acquire Zika virus from the environment, such as sewage containing the virus, and are able to transmit the virus to mammalian hosts (Du et al., 2019). Camptothecin, 1-hydroxycamptothecin, irinotecan and topotecan have been shown to inhibit Zika virus replication in human cells (Song et al., 2021). Therefore, a camptothecin-based anti-larval compound may also have a role to play in reducing viral transmission by acting on the virus in the insect.

### Safety for larvicidal use

Camptothecin and related compounds are cytotoxic to mammalian cells, which underpins their use in chemotherapy. We do not underestimate the challenge of deploying such compounds in the environment as larvicides. The target of camptothecin, topoisomerase I, is a highly conserved enzyme, with all the residues that contact the drug in a topotecan / human topoisomerase I crystal structure (Staker et al., 2002) conserved across insects (Fig 5). This likely limits our ability to make more insect-specific derivative molecules by exploiting differences in target binding.

**Fig. 5.**
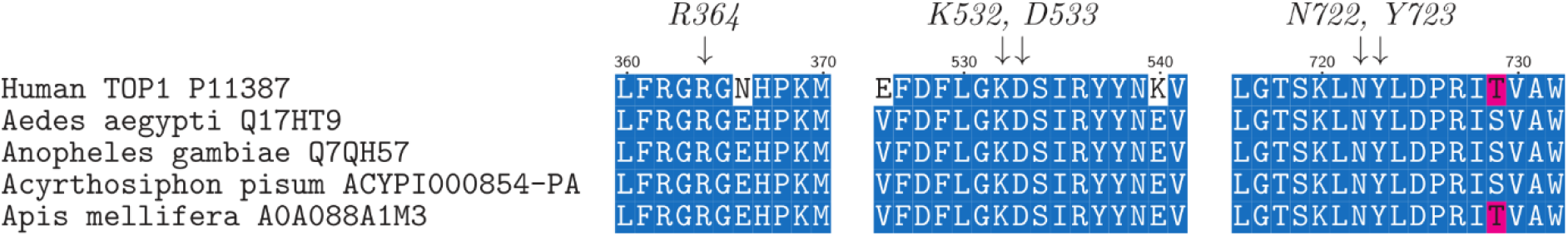
Conservation of the camptothecin binding site in topoisomerase I between insects and vertebrates. Residues of TOP1 that make direct contact with topotecan in a crystal structure (Staker et al., 2002) are highlighted above the alignment.

However, camptothecin itself has not found use in humans due to problems with pharmacokinetics. These problems include limited water solubility, rapid ring opening in plasma, where the active lactone is converted to an inactive carboxylate in plasma, and variable bioavailability, preventing oral dosing of camptothecin (Gupta et al., 2000; Herben et al., 1998). The route to clinical approval involved the development of water-soluble analogues of camptothecin, of which the first to be approved was topotecan, for intravenous administration, in 1996. Topotecan has appreciable oral bioavailability, around 35-40% (Herben et al., 1996), and ten-fold greater stability as the active lactone form in human blood compared to camptothecin (Burke and Bom, 2000). Topotecan was approved for oral administration in 2007. Interestingly, in our mosquito assay, topotecan was much less active than camptothecin. This suggests that there is potential for identification of camptothecin derivatives that have acceptable safety profiles by exploiting pharmacokinetic differences between target insects and people, such as drug access or metabolism. It should also be noted that organophosphates, currently important mosquito larvicides, have a poor safety record. Organophosphate poisoning, either due to occupational exposure or self-harm, kills an estimated 200,000 people each year (Eddleston and Chowdhury, 2016). This underscores the need to develop safer effective mosquito larvicides.

### Use as an adulticide

Females fed 100 μM and 10 μM camptothecin in a blood meal demonstrated significantly increased mortality (90% and 48.8% respectively) across the experiment compared to controls. Absolute blocking of egg laying was observed in females fed 100 μM camptothecin but no effect on egg laying, larval hatching or hatch percentage was observed for those fed 10 μM and many laid eggs prior to death during the experiment. This suggests that a concentration between 10 and 100 μM camptothecin would be required to impact the reproductive ability of *Ae. aegypti.*

TOP1 analogues are found in all eukaryotes and appear to be an essential enzyme during development in a wide variety of animals. During the process of DNA replication and transcription TOP1 is responsible for relaxing supercoiled DNA (Li et al., 2017). Knock outs of TOP1 are embryonically lethal in *Mus musculus* (Morham et al., 1996) and *Drosophila melanogaster* (Zhang, CX et al, 2000). TOP1 has been demonstrated to be essential for larval and pupal growth, oogenesis and embryogenesis in *D. melanogaster* (Zhang et al., 2000). Larvae are developing and undergoing more growth, cell replication and differentiation than in adults, which explains the greater susceptibility of larval stages that we observed.

Is it feasible to propose the use of a camptothecin derivative to target adult mosquitoes? We note the high concentrations of camptothecin (10 or 100 μM) that needed to be delivered in a blood meal to impact adult survival. Ivermectin is capable of killing *Anopheles* mosquitoes after they bite a human host who has taken the drug (Smit et al., 2019). However the concentration of ivermectin required for lethality is in the low nanomolar range (Dreyer et al., 2018), and it is well tolerated and is used widely for mass drug administration (MDA) control of helminths in areas where mosquitoes and malaria are a problem (Richards, 2017). We have already discussed the potential anti-viral use of camptothecin derivatives. Clearly compounds with improved potency against mosquitoes, as well as a much-improved safety profile, would need to be developed to be useful as anti-viral agents in humans with the additional benefit of controlling blood-feeding insect vectors.

## Acknowledgments

We thank Medicines for Malaria Venture for designing and supplying the Pandemic Response Box.

## Supporting information Captions

**S1 Movie. Recording of *Ae. Aegypti* larvae treated with camptothecin-derivatives at 100 μM, or the DMSO-only control, after 24 hours. Images were recorded every 100 ms.**

**S1 Table. Effect of all MMV Pandemic Box compounds on *Ae. aegypti* larval motility after 2 and 24 hours of treatment in the primary screen**

